# NOXA/MCL-1 axis determines cell-death decision between apoptosis and pyroptosis and the inflammatory secretome of breast cancer cells treated with anti-mitotics

**DOI:** 10.1101/2023.10.06.561231

**Authors:** Alison Dumont, Fabien Gautier, Quentin Batard, Catherine Guette, François Guillonneau, Mario Campone, Philippe Juin, Sophie Barillé-Nion

**Affiliations:** CRCI2NA, Inserm 1307, CNRS UMR 6075, Nantes Université, Université Angers, Equipe Labellisée LIGUE Contre le Cancer; ICO, Saint Herblain 44 France

## Abstract

Understanding how the malignant cells respond to chemotherapy is essential to prevent the development of resistance and to improve the efficiency of anti-cancer drugs. Recently, we established that, by intrinsic and paracrine mechanisms, taxol treatment in breast tumor cells increases NOXA a pro-apoptotic protein functioning as an endogenous inhibitor of survival protein MCL-1, thereby enhancing cytotoxic load on the compensatory survival protein BCL-xL. We herein sought to define the contribution of NOXA/MCL-1 to the modality of cell death secretome composition upon anti-mitotic treatment associated with a BCL-xL antagonist.

We observed that genetic inactivation of NOXA (enforcing MCL-1 pro-survival activity) in cancer cells not only delays their death when exposed to taxol in combination with the BCL-xL antagonist A1331852, but also alters its morphological characteristics with the apparition of features evoking pyroptosis. We identified the Caspase3-GSDME axis as regulating pyroptotic-like features suggesting that NOXA may act as a negative regulator of this cell death process (and MCL-1 as a positive regulator for it). Furthermore, comparative analysis of secretomes from the NOXA proficient or deficient cancer cells treated by taxol reveals variations in inflammatory cytokine production including those of IL-1β and IL-18.

Thus, our results show that anti-mitotic treatments are able to induce death by apoptosis and/or pyroptosis depending on BCL-2 family balance in breast cancer cells. Furthermore, NOXA/MCL-1 ratio appears to control the communication between these two types of cell death and their associated extracellular inflammatory signals in coordination with the pore-forming gasdermin GSDME.

## INTRODUCTION

The initial objective of chemotherapy was to eradicate tumor cells, operating under the assumption that inducing their demise would result in the complete elimination of tumors and the recovery of patients. However, recent advancements in biological research focused on tumor cell death have unveiled two critical insights. First, there exists a diversity of interconnected pathways leading to cell death. Second, cell death is not merely an endpoint; it carries valuable information for the tumor microenvironment. This information can either stimulate an anti-tumoral response or fail to do so. Dying cells release various signals into their microenvironment, including damage-associated molecular patterns (DAMP) or stress-associated molecular patterns (SAMP), depending on the specific cell death processes they undergo (Yatim 2017, Kroemer 2022). These signals, constitutive or inducible, in turn, lead to differential tumor responses (Yatim 2015, Legrand 2019). Importantly, these signals may trigger potent co-stimulatory signals, either supporting or inhibiting the intrinsic tumor response to anticancer therapy or the immune antitumor response. Consequently, it is now envisioned that the modalities of therapy-induced cell death may be harnessed to provoke an effective antitumoral immune response, involving T-cell immunity, referred to as immunogenic cell death (Kroemer 2022, Petroni 2021).

Cytotoxic chemotherapies traditionally target rapidly dividing cells, disrupting processes such as DNA replication or cell division, which induce acute stress in affected cells. In response, these cells attempt to repair the damage or, when overwhelmed, trigger self-destruct mechanisms. The outcome depends significantly on the intensity of acute stress and can lead to either cell survival or programmed cell death (Legrand 2019). The cellular response varies according to the type of stress and the cellular context, involving distinct molecular pathways and the release of information (Bedoui 2020, Barillé-Nion 2020). Beside well-characterized apoptotic pathways (Juin 2013), recent studies have identified and decoded pyroptosis as a cell death process initially engaged in immune cells during infection and relying on gasdermin (GSDM) family of proteins (Man 2017). GSDM proteins, mainly GSDMD and GSDME, are membrane pore forming effectors associated with the release of inflammatory cellular contents involved in regulation of essential innate immune processes and cancer cell response to chemotherapies (Shi 2015, Wang 2017).

Accumulating evidence points to cell death as a potent regulator of inflammation, even during cancer therapies. Initially, apoptotic cells, which break down into membrane-enveloped fragments, were considered immunologically silent due to their rapid clearance by phagocytes. In contrast, pyroptotic or necroptotic lytic cell death processes trigger an inflammatory response (Yatim 2015, Newton 2021). Notably, mitochondria play a central role in determining whether a cell lives or dies, by modulating metabolic and respiratory activities as well as orchestrating regulated cell death (reviewed in Marchi 2023). The integrity of the outer mitochondrial membrane is governed by the BCL2 family proteins that regulate its permeabilization (MOMP) (Juin, 2013) and by the release of cytochrome-c, which activates the caspase-9-dependent apoptosome, leading to the execution of the intrinsic apoptotic pathway. Additionally, when apoptotic caspase activities are inhibited, MOMP triggers proinflammatory signaling through the release of mitochondrial DNA, activating the cGAS/STING pathway, ultimately leading to the activation of the type I-IFN and NF-kB pathways (Rongvaux 2014, White 2014). Recent studies have also highlighted that antimitotic therapies activate the cGAS/STING pathway, in response to chemo-induced micronuclei (Zierhut 2019), in particular in breast cancer cells where a cascade serves as a catalyst for a proapoptotic secretome within tumors, enhancing the cytotoxic effects of the drug (Lohard 2020). Of interest, this signaling activation enhances the response to genotoxic treatments and immunotherapy (Schadt 2019). However, when chronic the intrinsic cGAS/STING activation may also promote metastasis (Bakhoum 2018).

Therefore, understanding how tumor cells respond to chemotherapy-induced stress and the information they release into their microenvironment is crucial for unraveling the complex interplay that affects tumor progression or elimination.

In this study, we shift our focus to explore the biological consequences of antimitotic therapy in breast cancers, with the aim of better characterizing the stress and information released during cell death triggered by this treatment. Our findings shed light on a shift in cell death mechanisms driven by NOXA and GSDME. These components play a pivotal role in determining whether the apoptotic or pyroptotic cell death pathways are activated. Our comprehensive analysis of soluble proteins released during cell death reveals that antimitotic treatment induces the release of pro-inflammatory IL1-related cytokines, specifically IL-1β and IL-18, in breast cancer cell lines and human primary breast tumors. Importantly, the release of these cytokines depends on the specific cell death pathway that was engaged during antimitotic treatment. Consequently, the manipulation of NOXA or GSDME functions holds the potential to redirect the immune response, enhancing its anti-tumoral activity.

## MATERIALS and METHODS

### Cells lines and reagents

MDA-MB-231, MDA-MB-468 cells lines were purchased from ATCC and cultured in Dulbecco’s Modified Eagle Medium (DMEM) (Gibco, Saint Aubin, France) supplemented with 2 mM glutamine and 1% penicillin/streptomycin (Gibco) and 5% Fetal Bovine Serum (FBS) (Eurobio, Courtaboeuf, France). MCL-1 overexpressing cell lines were established by viral infection with retroviruses containing vector coding for MCL1 (plvx). Puromycin was used to select for MCL-1 expressing cells at 1μg/ml. For the CRISPR Cas9-induced knock-out (KO) cell lines, single guide (sg) RNA sequences targeting human genes were designed using the CRISPR design tool (http://crispor.tefor.net). The guide sequences described in Table 1 (NOXA (PMAIP1), GSDME (DFNA5), GSDMD, CASP1, CASP3, BAX, BAK) were cloned in the plentiCRISPRV2 vector that was a gift from Feng Zhang (Addgene plasmid #52961). Cells were selected using 1 μg/ml puromycin and KO or KD were confirmed by immunoblot analysis.

**TABLE 1:**
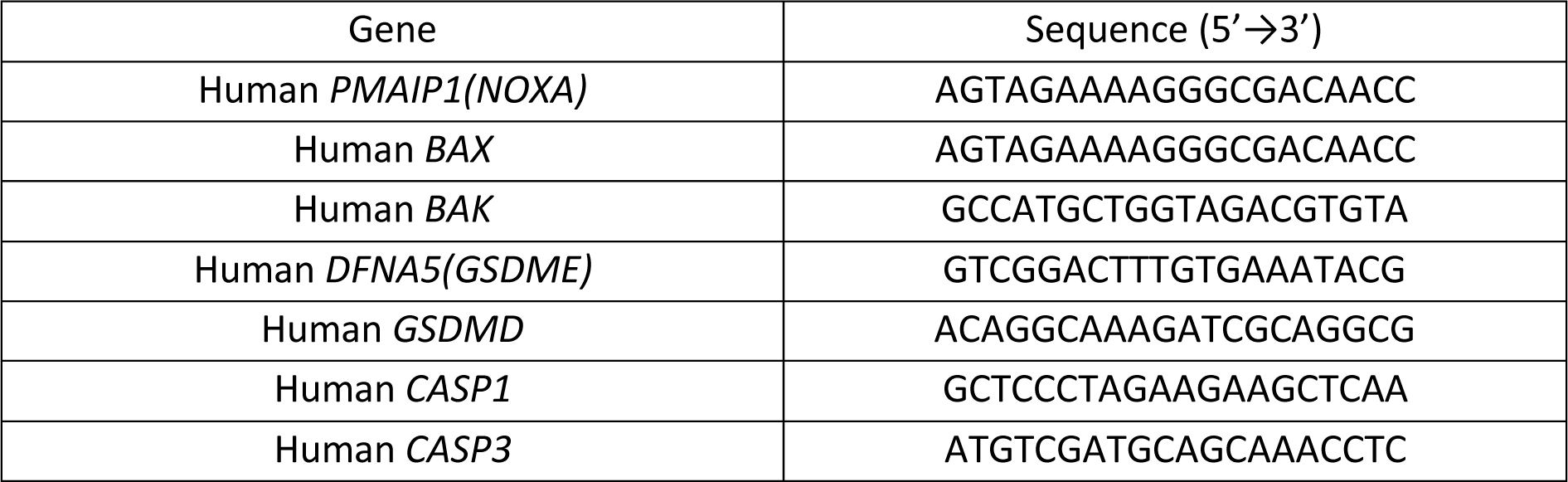
Guide sequences used in CRISPR-Cas9-based generation of KO cell lines.

For the preparation of conditioned media (CM), i.e. secretomes, 3.10^6^ cells were treated as indicated during 24h in 100mm plates, washed 3 times with PBS and cultured in DMEM without FBS for 48 additional hours. CM were then collected, centrifuged (2000 rpm,10 min) and concentrated using a polyethersulfone membrane to retain proteins above 3kDa.

In vitro treatments were used at the following concentrations: 100 nM A1331852 (*ApexBio, Houston, TX, USA*), 5 nM S63845 (Selleckchem, Houston, TX, USA), 70 nM taxol (Sigma-Aldrich, St Quentin Fallavier, France), 5 μM Q-VD-OPh (R&D Systems, Abindgon, UK).

### Incucyte imager-based videomicroscopy analysis

Cell death analysis was determined using an Incucyte Zoom imaging system (Sartorius). Cells were plated in 1mM thymidin-containing medium overnight for cell-cycle synchronization. Cells were then treated with taxol (70nM) or/and A1331852 (100nM) in presence of S63845 (500nM) or Q-VD-Oph (5μM) or not and AnnexinV dye (29011R-5UG, 50pg/µL), Caspase 3/7 dye (4440, Sartorius, 1/1000) and/or propidium Iodide (PI) 10 μg/mL (PI, P3566, Invitrogen) were then added. Plates were imaged every 60 or 120 min during 48h. The quantitative analysis was done using Incucyte cell-by-cell analysis software module (Sartorius). *n*=3 independent experiments; mean values ± SD. The qualitative analysis of cell death mode was evaluated on 30 dying cells in each condition, based on dye staining and on formation of blebs and/or a unique vacuole during cell death process. The two ways-ANOVA test was used for statistical analysis with GraphPad Prism 8.0 Software.

### Flow cytometry analysis

Apoptosis analysis was evaluated by staining cells with Annexin V-FITC and Propidium Iodide (PI) (130-092-52, Miltenyi Biotec, Bergisch Gladbach, Germany), according to manufacturer’s instructions. Cytochrome-c release was evaluated using the cyto-c antibody (560263, BD Biosciences) after fixation/permeabilisation using the Foxp3 transcription factor binding/permeabilization diluent and concentrate kit (Ebioscience, Thermofisher scientific)

Flow cytometry analysis was performed on FACS Accuri C6 plus (BD Biosciences). All experiments were repeated at least 3 times. The two ways-ANOVA test was used for statistical analysis with GraphPad Prism 8.0 Software.

### Biochemical assays

For immunoblot analysis, proteins were obtained by lysing cells with RIPA buffer (89901, Thermofisher scientific) or by concentrating cell supernatants (secretomes) 5X on centricon 3KDa, before separation on SDS-PAGE and transfer on nitrocellulose membranes. Membranes were then incubated with primary antibodies 1h at room temperature with the following antibodies used at following dilutions: Actin (MAB1501, EMD Millipore, 1/2000), NOXA (ab13654, Abcam, 1/500), GSDME (ab215191, Abcam, 1/1000), Caspase-3 (9662S, Cell Signaling, 1/1000), IL-18 (ab207323, Abcam, 1/500), GSDMD (HPA044487, Atlas Antibody, 1/1000), BAK (3814S, Cell Signaling Technology, 1/1000) BAX (5023S, Cell Signaling Technology, 1/1000), MCL-1 (94296S, Cell Signaling Technology, 1/1000). Then membranes were incubated with the appropriate secondary antibodies for 1 h at room temperature. Immobilon Forte Western HRP substrate kit (WBLUF0100, Millipore, Marne la Coquette, France) was used for immunoblot revelation on the Fusion FX + system (Veber).

For ELISA assays, levels of IL1β in cell supernatants were determined according to the manufacturer’s protocol (437015, Biolegend).

Caspase3/7 activity was quantified in cell lysates using the Kit Caspase-Glo® 3/7 Assay Systems (G8091, Promega).

All experiments were conducted 3 times in independent experiments and results are presented as mean values ± SD and the 2-way ANOVA statistical analysis was carried out.

### Proteomic analysis

The proteomics analysis was done in collaboration with the Prot’ICO Proteomics Mass Spectrometry Facility at ICO. Briefly, MDA-MB-468 CT^CRISPR^ or NOXA^CRISPR-/-^ cell pellets or corresponding concentrated supernatants were resuspended in 200µL of Rapigest SF (Waters), and dithiothreitol was added to a final concentration of 5mM (DTT, AppliChem). Samples were incubated in a thermo shaker at 95°C for one hour, and probe-sonication performed twice 30 seconds at 20% powerusing an ultrasonic processor (130W, 20 KHz, Bioblock Scientific). Subsequently, cysteine residues were alkylated by adding 200mM S-Methyl methanethiosulfonate (MMTS, igma-Aldrich) to a final concentration of 10mM (incubated at 37°C for 10min). Sequencing-grade trypsin (Absciex) was added in a ratio ≥2µg per 100µg of protein (incubated at 37°C overnight). The reaction was stopped with formic acid (9% final concentration) and incubated at 37°C for one hour, and the acid-treated samples were centrifuged at 16,000g for ten minutes. Salts were removed from the supernatant and collected in new reaction microtubes using self-packed C18 STAGE-tips.Peptide concentrations were finally determined with the Micro BCA™ Protein Assay Kit (Thermo Fisher Scientific). 200ng of each sample were analyzed by a LC-MS/MS system consisting in a nanoElute system (Bruker Daltonics GmbH) with an Aurora series reversed-phase C18 column (25 cm × 75 μm i.d., 1.6 μm C18, IonOpticks) heated to 50 °C in line with and coupled to a TimsTOF Pro2 (Bruker Daltonik GmbH). A gradient of 2–35% B, where mobile phase A was 0.1% formic acid in Milli-Q grade water and B was 0.1% formic acid in acetonitrile, was used for 60min. The total run times, including a ramp up to 35–95% B to clean the column and prep for the next sample was 90 min. Measurements were acquired in DIA-PASEF mode (data independent acquisition parallel accumulation serial fragmentation). The *m/z* range default was 400–1201 with an IM range of 0.6–1.43 1/K0 [V s/cm2], which corresponded to an estimated cycle time of 1.80 s. Mass spectrometry data were analyzed using Spectronaut^TM^ v16.0 (Biognosys AG). The direct DIA workflow in Spectronaut^TM^ was used for analyzing the DIA-PASEF dataset without a library from the DDA runs (library-free). TheUniprot curated Human protein database (release April 2022) served as template instead, with trypsin cleavage specificity, complete cysteins methylation and possible methionine oxidation. Otherwise default analysis settings were applied. Retention time prediction type was set to dynamic iRT, and calibration mode was set to automatic. These settings also included mutated decoy method, and cross run normalization was enabled. The FDR (false discovery rate) was estimated with the mProphet approach, and Q-value cut-off for both precursor and protein was set to 1%. Interference correction was enabled for quantification which required a minimum of 2 precursor ions and 3 fragment ions. The term protein refers to protein groups as determined by the algorithm implemented in Spectronaut^TM^. These data were further processed by dividing MS counts by the number of cells within each sample to standardize measurements to the cell level. Missing data were imputed using iterative PCA algorithm (missMDA package) and the factoMineR package was used to achieve PCA analysis. Statistical significance of differentially expressed proteins were assessed using t.test function (stats R package). The resulting protein lists were ranked according to log2FC values, and top50 proteins were passed to the EnrichR enrichment analysis tool ((https://maayanlab.cloud/Enrichr/). Volcano Plots were generated using the corresponding R package, and ggplot2 was used to produce boxplot figures.

### Transcriptomic analysis

For RTqPCR experiments, total RNA was isolated using Nucleospin RNA plus (Macherey Nagel, Hoerdt, France) and transcribed into cDNA by Maxima First Strand cDNA synthesis Kit (Thermo Scientific, Illkirch, France). Quantitative RT-PCR (qPCR) was performed using the EurobioGreen qPCR Mix Lo-Rox with qTOWER (Analityk-jena, Jena, Germany). Reaction was done in 10 μl final with 4 ng RNA equivalent of cDNA and 150 nM primers. Primers sequences used for DNA amplification are the following pairs:

*IL18*:ATTACTTTGGCAAGCTTGAATCTAAATTATCAGT/TACCTCTAGGCTGGCTATCTTTATACATACT

*IL1B:* TGGCAATGAGGATGACTTGT/GGAAAGAAGGTGCTCAGGTC

*ACTB:* AGAAAATCTGGCACCACACC/CAGAGGCGTACAGGGATAGC

RTqPCR experiments were conducted in 3 independent experiments and results are presented as mean values ± SD and the 2-way ANOVA statistical analysis was carried out.

For Nanostring analysis, gene expression was quantified with the NanoString nCounter platform in the nCounter Human (255 genes) Inflammatory Panel (NanoString Technologies) using 50ng of total RNA from patient-derived luminal tumor explants after a 48h ex vivo incubation with taxol 700nM or not (as described in Lohard et al). The code set was hybridized with the RNA overnight at 65°C. RNA transcripts were immobilized and counted using the NanoString nCounter Sprint. RCC files were read using the nanostring R package. Positive control normalization and housekeeping genes normalization were done according to Gene Expression Data Analysis Guideline instructions (https://nanostring.com/wp-content/uploads/Gene_Expression_Data_Analysis_Guidelines.pdf). Normalized data were then processed using standard R workflows.

### Statistical analysis

The two ways-ANOVA test was used for statistical analysis with GraphPad Prism 8.0 Software. Errors bars represent standard deviation (SD). The symbols correspond to a P-value inferior to *0.05, **0.01, ***0.001, ****0,0001

## RESULTS

### NOXA controls pyroptotic-like cell death decision in triple negative breast cancer cells in response to cytotoxic antimitotic chemotherapy

As we previously observed that NOXA protein played a major role in cell death decision induced by the combination of antimitotic and BCL-xL-targeting BH3 mimetic in triple negative breast cancer cells (Lohard 2020), we decided to explore to which extent it could delay cell death and impact its execution.

Using a videomicroscopic approach to track each individual cell response to treatments, we first analyzed single cell death morphology and its kinetics arising in the triple negative breast cancer cell lines MDA-MB-231 and MDA-MB-468 functionally deleted of NOXA by inactivating NOXA coding gene (PMAIP1) by CRISPR-Cas9 technology, (NOXA^CRISPR-/-^) or not (CT^CRISPR^), in response to taxol treatment combined to the BCL-XL specific BH3 mimetic A1331852. NOXA gene inactivation (Fig 1A, E) not only led to delayed cell death occurrence when cancer cells were exposed to the cytotoxic treatment (Fig 1B, F), but also completely shifted cell death mode towards a process characterized by the appearance of a ballooning morphology inherent to the formation of a unique cytoplasmic vacuole strongly evocating a pyroptotic-like process (Fig 1C, G), as illustrated in Fig 1D, H. Of note, we observed an intrinsic heterogeneity in the morphology of cell death response to cytotoxic treatment in control cells (CT^CRISPR^) that underwent a dying path resembling apoptosis (50%) or pyroptosis (50%) in MDA-MB-468 cells (Fig 1C, D) and an initial apoptotic-like process (characterized by blebbing initiation and formation of apoptotic corps) evolving in final ballooning process in the residual main corps, in MDA-MB-231 cells (Figure 1G, H).

**Figure 1:**
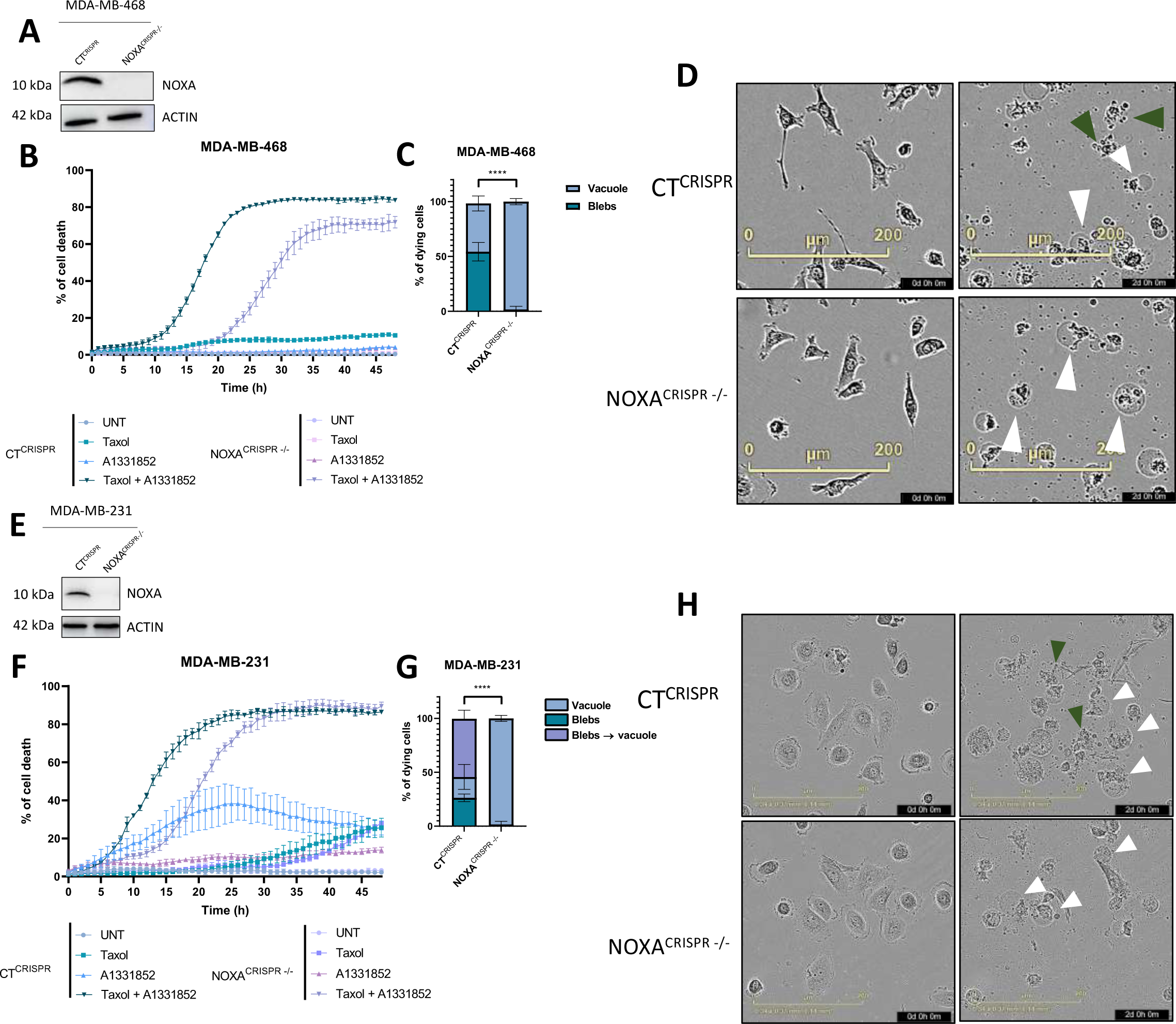
Morphological analysis of cell death process by videomicroscopy in CT^CRISPR^ and NOXA^CRISPR-/-^ cells. **A, E**: Immunoblot analysis of NOXA expression in CT^CRISPR^ and NOXA^CRISPR-/-^ MDA-MB-468 and MDA-MB-231 cells, respectively. **B, F**: Kinetics of single cell death analysis in untreated (UNT) or treated by taxol (70nm) or the BCL-xL inhibitor A1331852 (100nm) alone or combined during 48h, in CT^CRISPR^ and NOXA^CRISPR-/-^ MDA-MB-468 and MDA-MB-231 cells, respectively. **C, G**: % of cells dying either with blebs or with a unique vacuole after a 48h-exposure to combined treatment in CT^CRISPR^ and NOXA^CRISPR-/-^ MDA-MB-468 and MDA-MB-231 cells, respectively. **D, H**: Illustrative images at 0h and 48h of combined treatment with white arrows pointing vacuole-associated cell death and green arrows pointing blebs associated cell death in CT^CRISPR^ (upper panel) and NOXA^CRISPR-/-^ (lower panel) MDA-MB-468 and MDA-MB-231 cells, respectively. Data are mean ± SD of 3 independent experiments and statistical analysis were obtained using a two ways-ANOVA test.

### MCL-1 activity phenocopies NOXA depletion promoting pyroptosis-like onset versus apoptosis in dying cells

Importantly MCL-1 overexpression (Fig 2A, D) efficiently protected cells from death since analysis of single cell fates indicated that an average of 45% of the cells were still alive at the end of the experiments. In the same time, MCL-1 promoted pyroptotic-like cell death engagement in the 55% dying cells (Fig 2B, C, E, F), as did NOXA depletion. Conversely, targeting MCL-1 with its specific BH3-mimetic S63845, restored the ability of NOXA-depleted cells to undergo fast apoptosis at the expense of pyroptotic-like cell death, as did the control cells (Fig 2G to J). These results argue that the canonical function of NOXA, as an endogenous MCL-1 antagonist, determines the cell death switch between apoptotic- and pyroptotic-like modes observed during the cytotoxic treatment. In addition, they also suggest that MCL-1 protects cancer cells from apoptosis as expected, but not from pyroptotic death.

**Figure 2:**
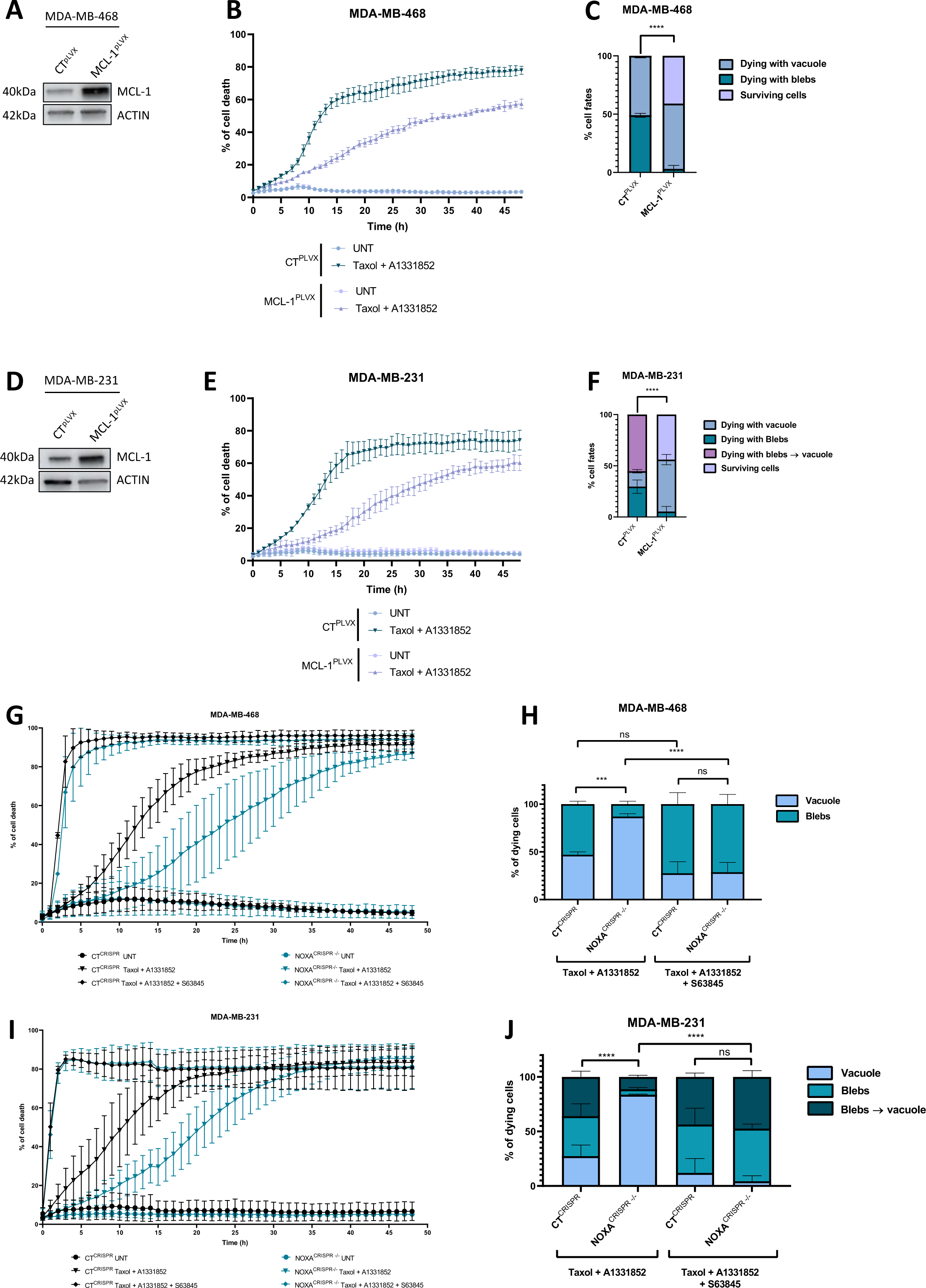
NOXA controls cell death process through MCL-1 inhibition. **A, D**: Immunoblot analysis of MCL-1 expression in CT^plvx^ and MCL-1^plvx^ MDA-MB-468 and MDA-MB-231 cells, respectively. **B, E**: Kinetics of single cell death analysis in untreated (UNT) or treated by taxol (70nm) and the BCL-xL inhibitor A1331852 (100nm) during 48h, in CT^plvx^ and MCL-1^plvx^ MDA-MB-468 and MDA-MB-231 cells, respectively. **C, F**: % of cell fates including dying with vacuole or blebs or surviving cells after a 48h-exposure to taxol and A1331852 in CT^plvx^ and MCL-1^plvx^ MDA-MB-468 and MDA-MB-231 cells, respectively. **G, I**: Kinetics of single cell death analysis, in untreated (UNT) or treated by taxol and A1331852 plus the MCL-1 inhibitor S63845 or not during 48h in CT^plvx^ and MCL-1^plvx^ MDA-MB-468 and MDA-MB-231 cells, respectively. **H, J :** % of dying cells with a vacuole or blebs or another cell death process after a 48h-exposure to taxol and A1331852 combined to S63845 or not, in CT^plvx^ and MCL-1^plvx^ MDA-MB-468 and MDA-MB-231 cells, respectively.

### Preferential pyroptotic-like cell fate relies on MOMP and CASP3 activity

Since NOXA/MCL-1 complexes participate in MOMP control, we first evaluated MOMP onset during the cell death process by first quantifying cytochrome-c (cyto-c) release in NOXA^CRISPR-/-^ cells upon treatment compared to CT^CRISPR^ after 36h of treatment corresponding to the complete cell death timing, in both cell lines. We could notice that NOXA gene deletion deeply limited cyto-c release compared to CT^CRISPR^ (Fig 3A, B, upper panels), while cell death rates were comparable (Fig 3A, B, lower panels). The importance of MOMP in the cell death process was further explored using double KO BAX/BAK MDA-MB-468 cells that were no more able to execute MOMP (Fig 3C). Videomicroscopy analysis indicated that in these cells, cell death was completely prevented (Fig 3D), highlighting the major role of MOMP in cell death process. Moreover, the pharmacological caspase-3 inhibition using the pan-caspase inhibitor Q-VD-Oph also stopped cell death process (Fig 3E) including the pyroptotic-like one. In the same way, deletion of CASP3 (CASP3^CRISPR-/-^cells) (Fig 3F) protected cells from cell death (Fig 3G, H) since they did not incorporate the viability exclusion dye, most of them however acquired a round condensed morphology at the end of the experiment, evocating a long-lasting mitotic arrest (Fig supp 1). Of note kinetics of caspase-3/7 activities detected during treatments, arose faster in CT^CRISPR^cells than in NOXA^CRISPR-/-^cells (Fig 3I, J), corresponding respectively to apoptotic-versus pyroptotic-like cell death in response to treatment. Altogether these results demonstrate that treatment-induced cell death either through apoptotic or pyroptotic-like process required MOMP onset and caspase-3 activity to take place. They suggest that pyroptosis-like process may happen under low MOMP, previously described as minority MOMP, when MCL-1, still operative in the absence of NOXA, efficiently mitigates the extent of MOMP per cell.

**Figure 3:**
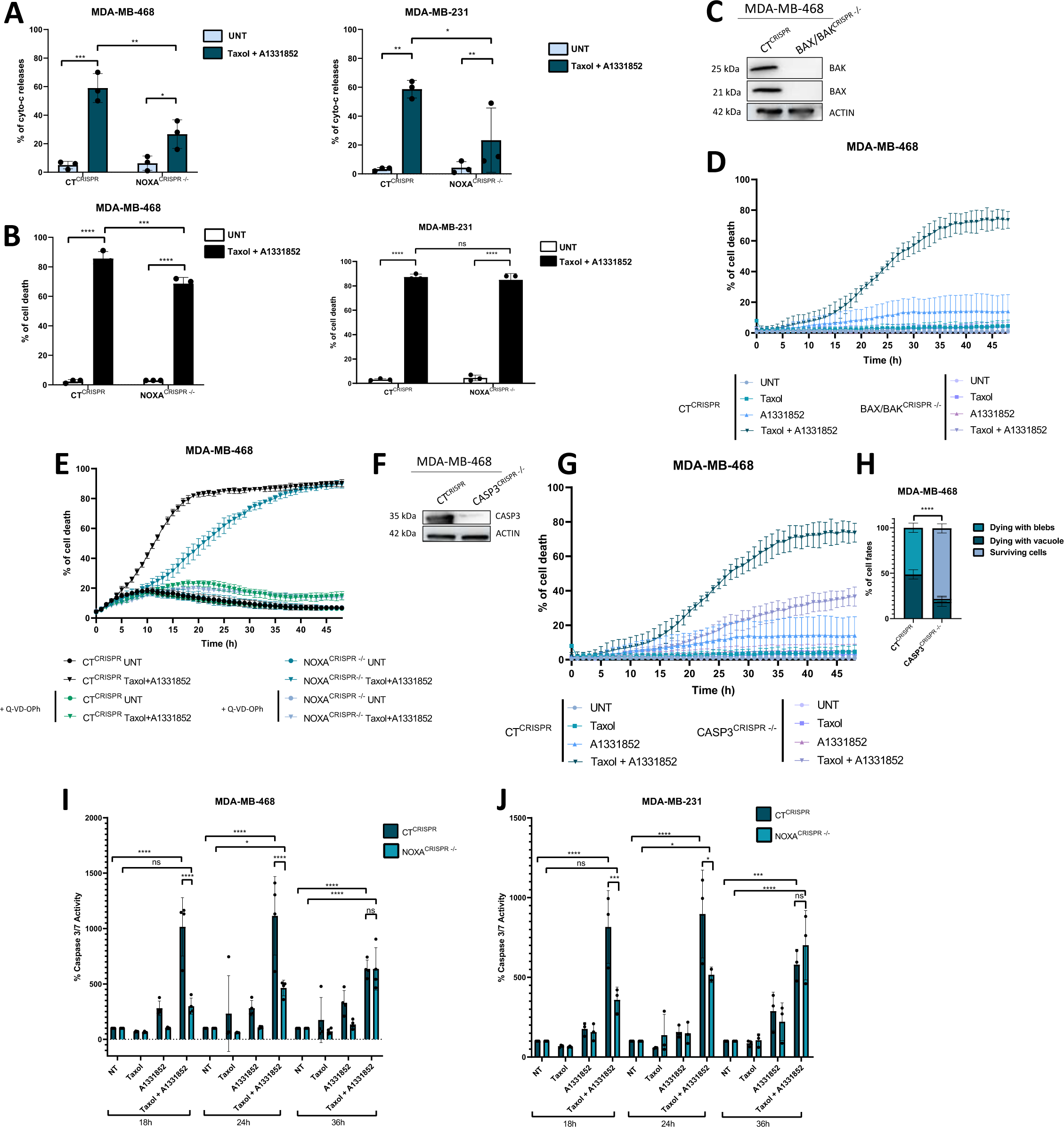
MOMP and casp3 activity are required for apoptotic and pyroptotic-like cell death course. **A, B**: Analysis of cyto-c release (upper panel) and cell death (lower panel) by flow cytometry in untreated (UNT) or treated by taxol or A1331852 alone or in combination during 36h, in CT^CRISPR^ and NOXA^CRISPR-/-^ MDA-MB-468 and MDA-MB-231 cells, respectively. **C**: Immunoblot analysis of BAX and BAK expression in CT^CRISPR^ and double KO BAX/BAK^CRISPR-/-^ MDA-MB-468. **D**: Kinetics of single cell death analysis, in untreated (UNT) or treated by taxol (70nm) or the BCL-xL inhibitor A1331852 (100nm) alone or in combination during 48h, in CT^CRISPR^ and double KO BAX/BAK^CRISPR-/-^ MDA-MB-468. **E**: Kinetic quantification of cell death during treatment by Taxol or A1331852 combination during 48h, in CT^CRISPR^ and NOXA^CRISPR-/-^ MDA-MB-468 in presence oF Q-VD-Oph or not. **F**: Immunoblot analysis of CASP3 expression in CT^CRISPR^ and CASP3^CRISPR-/-^ MDA-MB-468. **G:** Kinetics of single cell death analysis, in untreated (UNT) or treated by Taxol (70nm) or the BCL-xL inhibitor A1331852 (100nm) alone or combined during 48h, in CT^CRISPR^ and CASP3^CRISPR-/-^ MDA-MB-468. **H**: % of cells dying either with blebs, a unique vacuole or another cell death morphology after a 48h-exposure to combined treatment in CT^CRISPR^ and CASP3^CRISPR-/-^ MDA-MB-468. **I-J**: Kinetic quantification of Casp3/7 activity during treatment by Taxol or A1331852 alone or combined at 18, 24, 36 and 48h, in CT^CRISPR^ and NOXA^CRISPR-/-^ MDA-MB-468 and MDA-MB-231 cells.

### GSDME contributes to pyroptotic-like cell death onset in antimitotic-treated cancer cells

Since pyroptosis execution mainly relies on GSDM family, we first focused on GSDME that was significantly expressed in MDA-MB-231 and MDA-MB-468 cells (Fig 4A, E). GSDME gene depletion by CRISPR-Cas9 had no significant impact on cell death kinetics (Fig 4B, F), however it mostly prevented pyroptotic-like cell death onset in cancer cells upon cytotoxic treatment, to the benefit of enhanced apoptotic cell death rates (Fig 4C, H), as illustrated in Fig 4D, H. Importantly, a cleaved form of GSDME protein could be detected by immunoblot in treated cancers cells, compatible with its N-ter pore-forming fragment (Fig 4I, J). This cleavage in addition coincided with detection of activated caspase-3. Moreover, inhibition of apoptotic caspases using Q-VD-Oph potently prevented the cleavage of GSDME (Fig 4K, L). These results thus revealed that GSDME mainly contributes to treatment-induced pyroptotic-like cell death in breast cancer cells and that its activation by apoptotic caspases is necessary to its cleavage.

**Figure 4:**
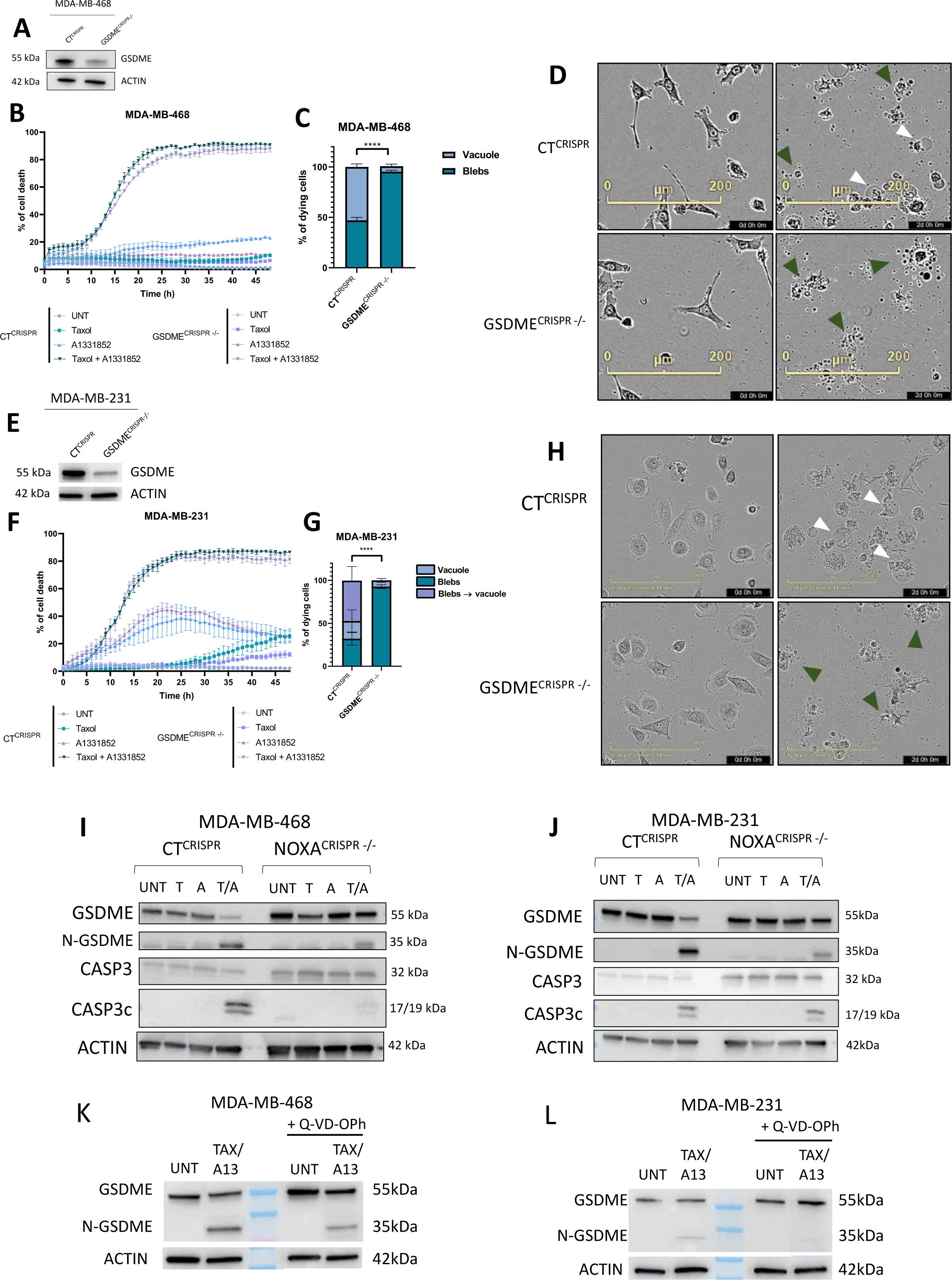
GSDME controls pyroptotic-like cell death induced by antimitotic-based chemotherapy, in correlation with its proteolytic cleavage. **A, E**: Immunoblot analysis of GSDME expression in CT^CRISPR^ and GSDME^CRISPR-/-^ MDA-MB-468 and MDA-MB-231 cells, respectively. **B, F**: Kinetics of single cell death analysis in untreated (UNT) or treated by taxol (70nm) or the BCL-xL inhibitor A1331852 (100nm) alone or in combination during 48h, in CT^CRISPR^ and NOXA^CRISPR-/-^ MDA-MB-468 and MDA-MB-231 cells, respectively. **C, G**: % of cells dying either with blebs or with a unique vacuole after a 48h-exposure to combined treatment in CT^CRISPR^ and GSDME^CRISPR-/-^ MDA-MB-468 and MDA-MB-231 cells, respectively. **D, H**: Illustrative images at 0h and 48h of combined treatment with white arrows pointing vacuole-associated cell death and green arrows pointing blebs associated cell death in CT^CRISPR^ (upper panel) and GSDME^CRISPR-/-^ (lower panel) MDA-MB-468 and MDA-MB-231 cells, respectively. **I, J**: Immunoblot analysis of GSDME, Casp3 and NOXA expression in CT^CRISPR^ and NOXA^CRISPR-/-^ MDA-MB-468 and MDA-MB-231 cells, respectively. **K, L**: Immunoblot analysis of GSDME cleavage upon treatment in presence of pancaspase inhibitor Q-VD-Oph or not, in MDA-MB-468 and MDA-MB-231 cells, respectively.

### Pyroptotic prone cancer cells preferentially produce immune regulatory secretomes

As dying cells may release specific factors depending on cell death process triggered by the therapeutic insult, we first conducted a comprehensive study of soluble protein factors produced (i.e. secretomes) by taxol-treated cells that are prone to trigger a pyroptotic cell death (using NOXA^CRISPR-/-^ cells) compared to CT cells from MDA-MB-468 cell line, using MS SWATH proteomics approach (LC-MS/MS proteomics analysis). This analysis included 4 replicates and a step of normalization by cell numbers in each sample was applied. Based on p-adjusted values less than 0.05, 2618 proteins were identified as being significantly differentially expressed in secretomes derived from taxol-treated NOXA^CRISPR-/-^ cells versus taxol-treated CT^CRISPR^, as illustrated in the volcanoplot presented in Fig 5A. Among them, 141 proteins were found to be decreased (log2(Fold Change) inferior to -0,5 or superior to +1), while 889 proteins were found to be increased. Gene enrichment analyses applied to proteomics, highlighted that differentially expressed proteins associated with immune signaling signatures including neutrophile degranulation or immune system, specifically in NOXA^CRISPR-/-^ cells that execute a pyroptotic program upon taxol-based cytotoxic treatment in comparison to CT ^CRISPR-/-^ cells (Fig 5B). When focused on cytokine /interleukin components, the analysis revealed that the production of the IL-1 related cytokine IL-18, or IL-11 and CXCL-1 relied on a NOXA-dependent regulation after taxol treatment (Fig 5C). The analysis also pointed out the absence of significant difference in IL-1α release between taxol-treated NOXA^CRISPR-/-^ cells and CT cells and surprisingly the absence of detection of IL-1β production. Of note, specific variations in complement system regulatory proteins including CD46, CD47, CD55 and CD59 or Annexin-A phospholipid-binding protein family ANXA1 and ANXA2, as well as the High mobility group box-1 (*HMGB1*), a representative damage-associated molecular patterns (DAMPs), were more present in taxol-treated NOXA^CRISPR−/−^ secretome samples than in CT^CRISPR^ ones (Fig 5D). In addition, regarding LDH considered as a pyroptotic marker, an increase in LDHA was detected in the secretome produced by NOXA^CRISPR-/-^ cells compared to CT^CRISPR^ cells (Fig supp 2A). Interestingly, marks of type I IFN dependent inflammatory process such as IFIT3, IFI30 and IFI35 were more produced in samples from NOXA deficient cells compared to CT^CRISPR^ ones after taxol treatment (Fig supp 2B), in accordance with our previous results in Lohard 2020 showing cGAS/STING/type I-IFN activation in taxol treated cells.

**Figure 5.**
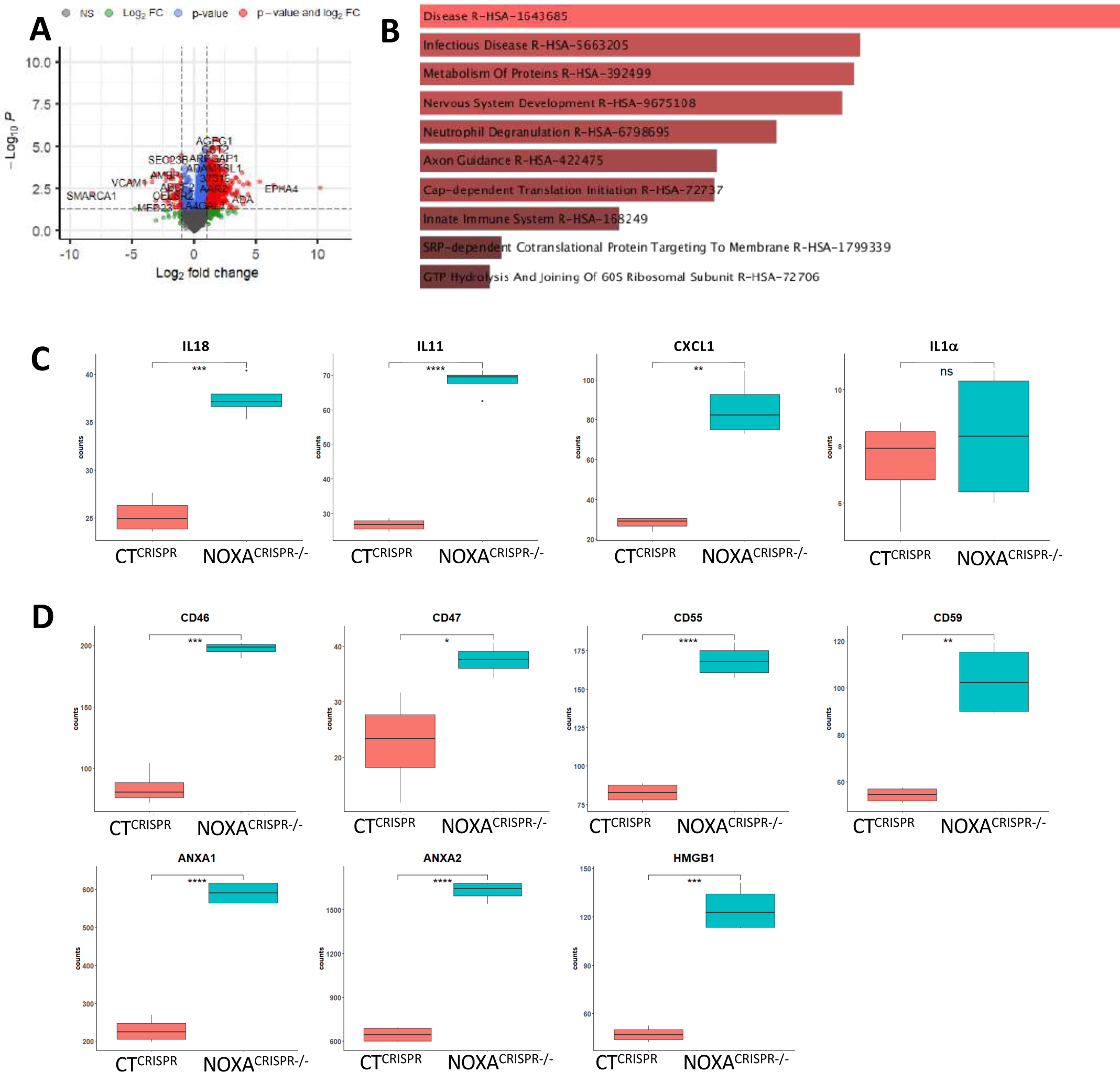
Proteomic analysis of secretomes from CT^CRISPR^ and NOXA^CRISPR-/-^ MDA-MB-468 after taxol treatment. **A:** Volcano plot showing overlap of proteins detected in the secretomes of CT^CRISPR^ and NOXA^CRISPR-/-^ MDA-MB-468 after taxol treatment. **B**: Gene ontology (GO) analysis of the unique 833 up-regulated proteins identified in the secretomes of taxol-treated NOXA^CRISPR-/-^ versus taxol-treated CT^CRISPR^ MDA-MB-468 fractions (log2FC >1 and False Discovery Rate (FDR)< 0.05), with the top 10 terms for the reactome plotted. **C, D**: Box plots of corresponding proteins identified in CT^CRISPR^ and NOXA^CRISPR-/-^ MDA-MB-468 secretomes after taxol treatment. Proteins are plotted according to their –log10P values and log2 fold enrichment.

Taken together, these results indicate that cancer cells prone to die by pyroptotic-like process upon antimitotic insult (i.e. NOXA^CRISPR−/−^), release more immune active factors than apoptotic prone cells, suggesting that cell death process may impact immune anticancer response.

### Taxol treatment triggered IL-1β and IL-18 secretion through GSDME complex process

Since we detected immune related signatures in secretomes from NOXA^CRISPR-/-^ treated cells, we further specifically explored the production of IL-1β (as a main indicator of pyroptotic process) and the IL-1-related interleukin IL-18, by dying cells. We first quantified IL-1β in cell culture supernatants by ELISA and we evidenced that taxol treatment significantly increased its production in NOXA^CRISPR−/−^ (Fig 6A). Importantly, IL-1β secretion was completely blocked in the absence of GSDME, even when cells were treated by taxol. Interestingly, this correlated with cell death modes preferentially triggered by taxol treatment, i.e pyroptosis proficient (NOXA^CRISPR-/-^) cells producing the highest amounts of IL-1β after taxol treatment.

**Figure 6.**
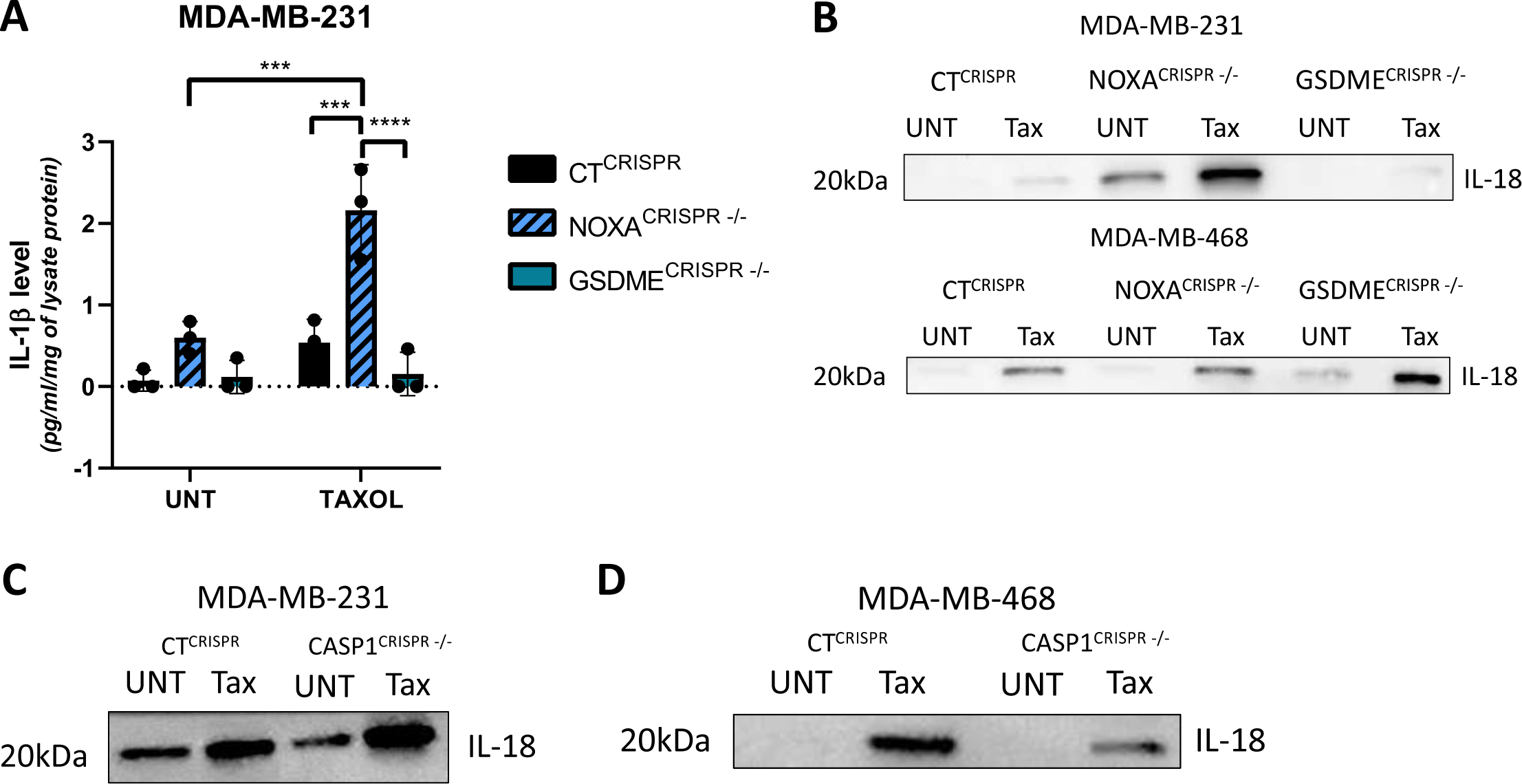
Taxol induced a NOXA/GSDME-dependent IL-1β secretion and a cCASP1-independent IL-18 secretion. **A**: IL-1β secretion meassured by ELISA in CT^CRISPR^, NOXA^CRISPR-/-^ and GSDME^CRISPR-/-^ MDA-MB-MDA-MB-231 cells after taxol treatment or not. **B**: Immunoblot analysis of IL-18 secretion in CT^CRISPR^, NOXA^CRISPR-/-^ and GSDME^CRISPR-/-^ MDA-MB-231 (upper panel) and MDA-MB-468 (lower panel) cells after taxol treatment or not. **C, D**: Immunoblot analysis of IL-18 secretion in CT^CRISPR^, and CASP1^CRISPR-/-^ MDA-MB-231 and MDA-MB-468 cells, respectively.

We further analyzed secretion of IL-18 in taxol-treated compared to untreated pyroptotic or apoptotic prone cells, using immunoblot analysis. We observed that IL-18 could be detected in its short form of 20 kDa corresponding to its mature form resulting from its intracellular proteolytic cleavage (Fig 6B). Importantly, IL-18 secretion was induced by taxol treatment in both CT^CRISPR^ and NOXA ^CRISPR−/−^ cell lines. However, IL-18 secretion was blocked in GSDME ^CRISPR-/-^ MDA-MB-231 cells thus revealing a prominent GSDME-dependent process in taxol-induced IL-18 secretion in these cells (Fig 6B, upper panel). In contrast, in MDA-MB-468 cells IL-18 production still occurred in the absence of GSDME suggesting an alternative secretion process (Fig 6B, lower panel).

To better understand how IL-18 was activated before secretion, we focused our attention on the canonical inflammasome pathway involving caspase-1 activity. We thus produced CASP1^CRISPR-/-^ cells to analyze their ability to secrete IL-18. We could not evidence significant change in IL-18 maturation or secretion between CASP1^CRISPR-/-^cells and CT^CRISPR^ cells (Fig 6C, D) arguing against a CASP-1 dependent pathway in IL-18 production upon taxol treatment. In addition, CASP1 deletion had no significant impact on cell death process (Fig supp 3A to D).

### Transcription of Inflammation-associated *IL1A/B, IL18 and TNF* genes is regulated by antimitotic treatment in breast cancer cells and tumors

Since IL-1β and IL-18 secretions were modulated by taxol treatment and/or cell death process, we performed qPCR analysis to evaluate the transcriptional activity of their corresponding genes. *IL-1B* gene transcription was strongly induced by taxol in CT ^CRISPR^ cells MDA-MB-231, as were those of *IL1A* and *TNF* (Fig 7A). In contrast *IL18* coding mRNAs were not induced by taxol treatment in these cells but even repressed (Fig 7A).

**Figure 7.**
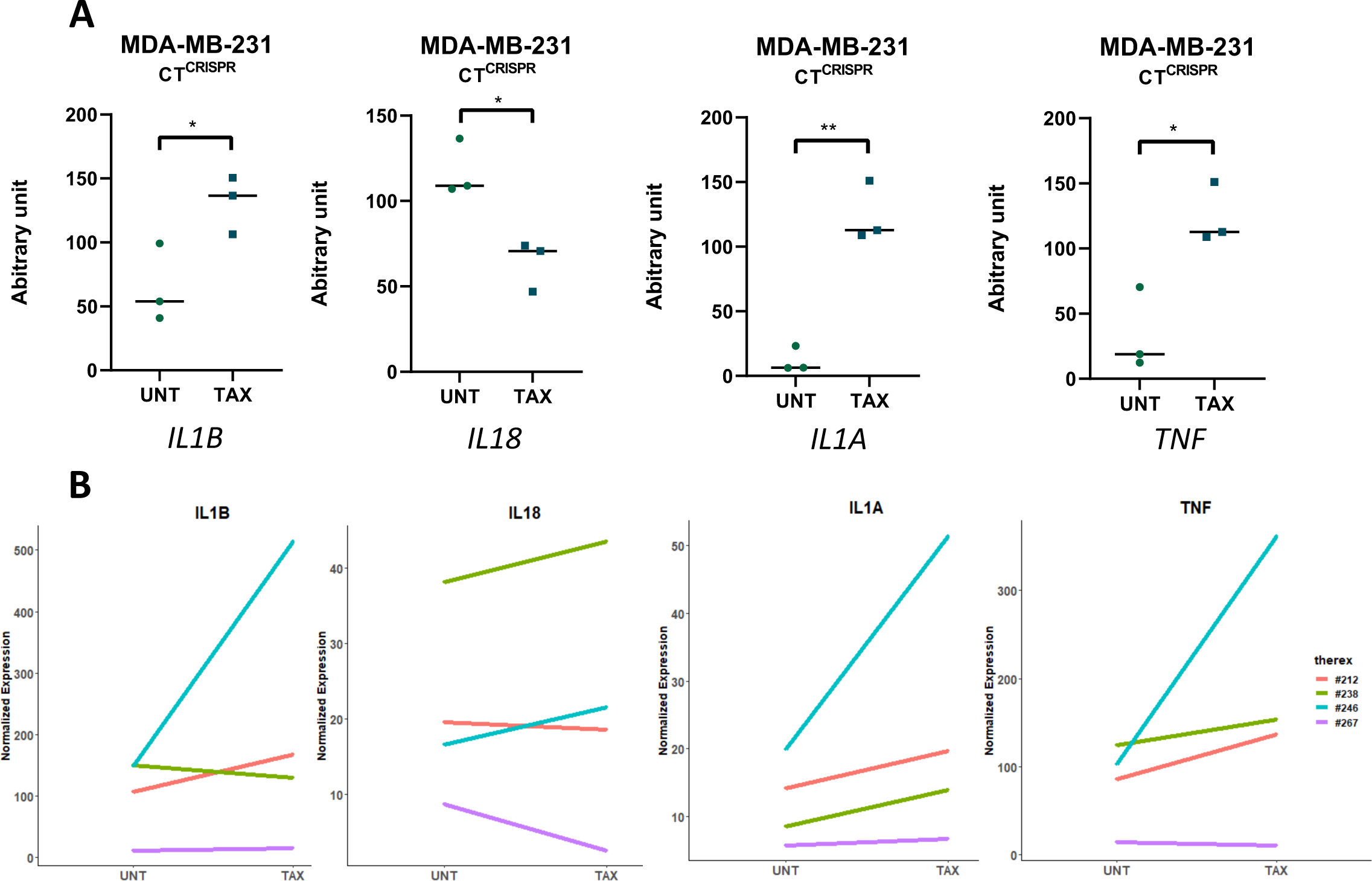
Taxol treatment induced IL1A/B but not IL18 gene transcription in breast cancer cell lines and human primary breast tumors. **A**: Q-PCR analysis of *IL1A/B, IL18* and *TNF* mRNA in CT^CRISPR^ MDA-MB-231 cells after taxol treatment or not. **B**: Transcriptomic analysis (*IL1A, IL1B, TNF and IL18*) using the Nanostring Inflammatory genes’ panel after a 48h ex vivo treatment with taxol (TAX) or not (NT) in 4 human primary breast tumors, among them 3 taxol-sensible (red, green and blue) and 1 taxol-resistant (purple).

We further interrogated these points in primary tumors, using Nanostring technology approach. We compared inflammation gene expression in 4 primary tumors from patients, after a 48h in vitro incubation with taxol or not (as described in Lohard 2020). The expressions of *IL1B* or *IL1A* genes were increased upon taxol treatment in 3 samples (Fig 7B). In contrast *IL18* gene expression was not or slightly induced in the taxol-treated tumors. Of interest, *TNF* gene expression was induced in 3 of the tumors, coinciding with our previous results reporting TNF activation in taxol-treated breast cancer tumors. We could notice that in this panel, the taxol-resistant tumor (that is the one in which taxol treatment induced the less active caspase-3 in tumor cells) showed no *TNF* gene regulation, the weakest regulation of *IL1B and IL1A* and a decreased *IL18* transcription upon taxol treatment.

Altogether, these results indicate that taxol triggers the production of IL-1β, which is linked to an upregulation in IL1B gene transcription while the increased IL-18 secretion, also observed following taxol treatment, does not coincid with the induction of its gene transcription. Furthermore, this phenomenon relies on GSDME-dependent or independent pathways depending on breast cancer cells.

## DISCUSSION

Cell death is a complex process, involving various factors that can tip the delicate balance between life and death, as well as between pro- and anti-inflammatory signals within the microenvironment. In our study, we demonstrate that antimitotic therapy in breast cancer cells triggers a specific type of cell death, which follows either an apoptotic or pyroptotic pathway depending on the intrinsic expression of NOXA and GSDME. This choice of pathway is crucial because it determines the release of inflammatory signals in the microenvironment. Importantly, we found that the absence of NOXA not only delays cell death in breast cancer treated with antimitotic taxol, but it also promotes pyroptosis over apoptosis. The major role of NOXA in response to anticancer drugs relies on its antagonist activity on MCL-1 (Dumont 2020). Our results indeed reveal an unexpected role of MCL-1 in controlling the crosstalk between cell death pathways induced by chemotherapy in cancer cells. In addition to its initial implication in genotoxic-induced cell death as a p53-target gene (Oda 2000), NOXA also supports cancer cell death triggered by antimitotics and proteasome inhibitors (Lohard 2020, Qin 2005). Of note, destabilization of its mRNA by oncogenic driver-targeting therapies such as BRAF or EGFR inhibitors, has been reported as an adaptation mechanism leading to dependence on MCL-1 in treated cancer cells and drug resistance (Montero 2019). In the same line, analysis of post-neoadjuvant chemotherapy residual disease in triple-negative breast cancer (TNBC), evidenced amplification of *MCL1* gene as one of the most prevalent genomic alteration in these resistant tumors (Balko 2014). Altogether, these data highlight NOXA and MCL-1 as major regulators of response to anticancer treatments.

We further dissected the molecular mechanisms underlying this effect and found that the timing of events such as mitochondrial outer membrane permeabilization (MOMP), cytochrome-c release, and caspase-3 activity plays a decisive role in determining whether pyroptosis or apoptosis commitment occurs. The onset of MOMP is of utmost importance as its inhibition through the use of BAX/BAK double KO cells, entirely prevented cell demise via either apoptosis or pyroptosis. Additionally, enforced expression of MCL-1 which protects from MOMP, efficiently curtailed cell death by apoptosis, however some cells still succumb to pyroptosis. This, coupled to reduced cyto-c release and caspase-3 activity, observed in the absence of NOXA, suggests the possibility of a “sub-apoptotic MOMP”, inadequate to trigger apoptosis but sufficient to foster pyroptosis. Tait and colleagues reported that an incomplete MOMP that spared intact mitochondria was compatible with the recovery of a new viable mitochondria network necessary to cell survival when apoptotic caspases were blocked (Tait 2010). This limited MOMP may also participate in long term resistance and aggressivity in cancer cells, in relation to DNA damage increase in these cells relying on increased activity of the Caspase-activated DNAse (CAD) and on complex regulation of mitochondrial dynamics (Ichim 2015, Cao 2022). Conversely, blocking caspase activity during MOMP has also potent anti-tumorigenic effects, mainly through NF-kB pathway control of necroptotic cancer cell death and immune responses (Giampazolias 2017). Our results support the concept that the sub-optimal MOMP induced by chemotherapy may predispose cancer cells to engage in pyroptosis, since mild MOMP was sufficient to engage cancer cell towards pyroptosis rather than apoptosis. They also suggest that GSDME may provide a failsafe mechanism against the accumulation of genetically instable (apoptotic resistant) cells and a contrario its downregulation may contribute to genetic instability. It’s worth mentioning that aggressive drug-tolerant cancer cells resulting from sub-lethal treatments, have been shown to exhibit increased susceptibility to ferroptosis (Kalkavan 2022). Thus, comprehending how cancer cells precisely respond to chemo-induced stress is of paramount importance to anticipate the emergence of resistance and to identify new therapeutic opportunities.

We further demonstrate that the expression of GSDME decisively determines the nature of cell death observed, with GSDME emerging as a key protagonist in pyroptotic cell death induced by chemotherapy. Recent research has identified gasdermins as primary executors of pyroptosis, forming membrane pores during caspase-1 activation and regulating essential innate immune processes (Shi 2017). In addition to its potential role as a tumor suppressor gene in breast tumors, due to its ability to boost antitumor immune response (Zhang 2020), recent findings suggest that GSDME can facilitate the progression of cell death toward pyroptosis in chemo-treated cancer cells, relying on caspase3 mediated and membrane pore-forming processes (Wang 2017, Rogers 2019). Our results substantiate these observations by highlighting the critical role of caspase-3 in initiating pyroptotic cell death and the proteolytic cleavage of GSDME into fragments capable of forming membrane pores.

Many regulated cell death processes hinge on the activation and recruitment of pore forming proteins that serve as executioners of specific cell death pathways: BAX/BAK in apoptosis, GSDM in pyroptosis or MLKL in necroptosis, using specific pore architectures (Vandenabeele 2023). Recent research also reported the role of the cell surface NINJ-1 protein in mediating the plasma membrane rupture (PMR) during cell death processes and the release of LDH (a standard measure of PMR) or DAMPs including HMGB1, in macrophages (Kayagaki 2021). In the opposite endosomal sorting complexes required for transport (ESCRT) can repair small wounds and pores in the plasma membrane, limiting IL-1β release in particular GSDMD-dependent pyroptosis in inflammasome-activated cells (Rühl 2018). The possible roles of these proteins in regulating the plasma membrane rupture in cancer cell death remain unanswered questions.

Given that apoptosis and pyroptosis are often considered as opposite processes in inflammation, we conducted a extensive proteomic analysis to compare the proteins released by antimitotic-treated cells predisposed to undergo either apoptosis or pyroptosis, based on the presence or absence of NOXA or GSDME. Our analysis clearly reveals that cells undergoing pyroptosis in response to antimitotic treatment release a more proinflammatory secretome compared to apoptotic cells. This is underscored by the increased levels of inflammatory cytokines such as IL-1b, IL-11, IL-18, and CXCL-1, along with various components of the complement pathway and the DAMP HMGB1. Additionally, we observed the presence of type I-IFN markers like IFIT3, IFI30, and IFI35 in these secretomes, aligning with our prior findings of cGAS-STING-dependent IFN-I activation in antimitotic-treated breast cancer cells (Lohard 2020).

Turning our attention to IL-1β and IL-18, we found that antimitotic taxol induced IL-1β production specifically during pyroptosis compared to apoptosis. Furthermore, our results shed light on a GSDME-dependent mechanism for IL-1β release during chemo-induced pyroptosis, as evidenced by the significant reduction of IL-1β production upon GSDME depletion. This aligns with previous research demonstrating GSDME’s role in enabling IL1β release, particularly in macrophages (Zhou 2021, Afonina 2015). On the other hand, with regard to IL-18, our findings indicate that antimitotic treatment leads to increased intrinsic IL-18 secretion in breast cancer cells. However the involvement of GSDME in this secretion appears to vary depending on the specific breast cancer cell lines. Notably, it is indispensable in MDA-MB-231 cells but dispensable in MDA-MB-468 cells. Intriguingly, we observed no discernable functional role of caspase-1 in IL-18 production, as its depletion had no discernable effect. It’s worth noting that the secretion of inflammatory cytokines may utilize multiple secretion pathways contingent upon the cellular context (Rébé 2020). Further research is warranted to unravel the mechanisms governing IL-18 production in the extracellular compartment and its intracellular maturation, including the identification of specific caspases or proteases contributing to its cleavage during chemo-induced pyroptosis.

Of utmost importance, we observed a concomitant increase of gene transcription levels of *IL1B* and *IL1A* in response to antimitotic taxol, a finding evident on the breast cancer cell line MDA-MB-231 and corroborated by three patient-derived tumors exposed to taxol ex vivo (Lohard 2020), corresponding to increased secretion of IL-1β. It is noteworthy that IL-1β exhibits both positive and negative roles in cancer, influencing immune cells with either pro- or anti-tumoral effects during oncogenesis or cancer treatments. In particular, high mRNA expression levels of IL1B have been associated with better prognosis in breast cancer patients, notably in TNBC, where treatment with anti-IL-1b antibodies in preclinical models, facilitated adaptive antitumor cell immunity and potentiated the effects of immune checkpoint inhibitors (Kaplanov 2019). Alternatively, activation of inflammasome using the omega3 fatty acid DHA, or the natural alkaloid Barberin, has yielded similar encouraging results (Sonnessa 2020). These promising findings open up new therapeutic avenues to enhance the treatment response in TNBC.

Remarkably, antimitotic treatment did not induce *IL18* transcription in breast cancer cells lines, despite a clear increase in secretion. This leads us to postulate that taxol may stimulate a specific production of IL-18 that is already present within the cells, a process warranting further investigation. Additionally, taxol had limited effects on *IL18* gene transcription in patient-derived tumors. Notably, the taxol-resistant tumor exhibited the lowest cytokine-related gene transcription, including TNF. However, it is important to acknowledge that we were unable to differentiate *IL18* expression by malignant or non-malignant cells within patient tumors. Nonetheless, our overall findings suggest a complex regulation of IL-18 secretion in response to antimitotic agents in breast cancer cells and tumors, implicating multiple levels, including gene transcription, intracellular proteolytic maturation and secretion. Given that IL-18 possesses diverse inflammatory activities and is known to modulate immune responses, especially by enhancing Th1 proliferation and NK cell activation (Molgora 2018, Park 2017), which can in turn impact their antitumor activities, it becomes imperative to conduct further research to delineate whether targeting IL-18 would prove beneficial or detrimental in cancer therapy.

In summary, our data underscores the profound influence of cell death processes induced by chemotherapy on the release of proinflammatory factors. Given the growing interest in inducing more immunogenic cell death during chemotherapy, it becomes imperative to design thoughtful combinations of chemotherapy and immunotherapy to maximize their synergistic potential. The promising outcomes observed in breast cancer with the combination of immune checkpoint inhibitors with chemotherapy in a neoadjuvant setting (Bardia 2021, Shahbandi 2022), underscore the importance of understanding the processes that govern the production of these proinflammatory factors, given their capacity to modulate the anti-tumor immune response. Our results advocate for the promotion of pyroptosis over apoptosis in response to cytotoxic treatment through intervention on the NOXA/MCL-1 axis, as this may offer stronger support for modulating the anticancer immune response. Nonetheless, a comprehensive view of the interplay between IL-1β and IL-18 and their effects on the immune response now warrants further investigation.

## ACKNOWLEDGEMENTS

We thank the Ligue contre le Cancer Grand Ouest for supporting XXX. We thank Aurélie Fétiveau for their technical support to produce KO contructs and Laurent Maillet and Florian Chocteau for fruitful discussion. We thank the CytoCell facility (SFR-Santé F Bonamy UMS016) for technical support.

## AUTHORS CONTRIBUTION

AD, PPJ and SBN conceived the study and designed the work. AD, QB, SBN performed experiments. AD, FG, PPJ and SBN contributed to data analysis. CG and FG contributed to proteomic analysis. MC provided tumor samples. PPJ and SBN obtained fundings and SBN wrote the paper.

## ADDITIONAL INFORMATION

Correspondence and requests for materials should be addressed to SBN.

## LEGENDS TO SUPPLEMENTARY FIGURES

**Supplementary figure 1:**
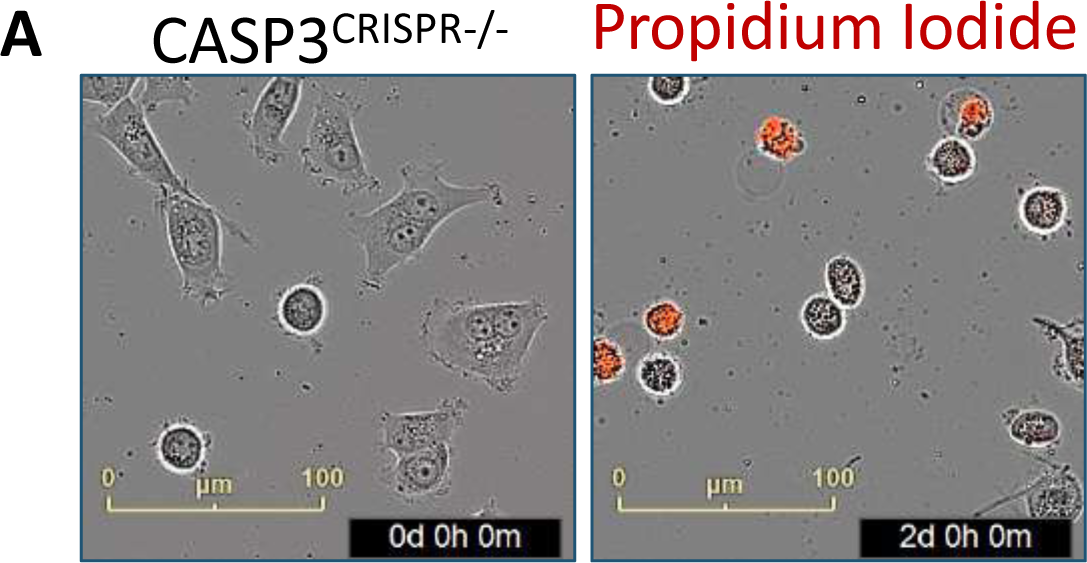
CASP3^CRISPR-/-^ MDA-MB-468 cells partially died upon treatment. Illustrative images at 0h and 48h of combined treatment in CASP3^CRISPR-/-^ MDA-MB-468, in presence of propidium iodide.

**Supplementary figure 2:**
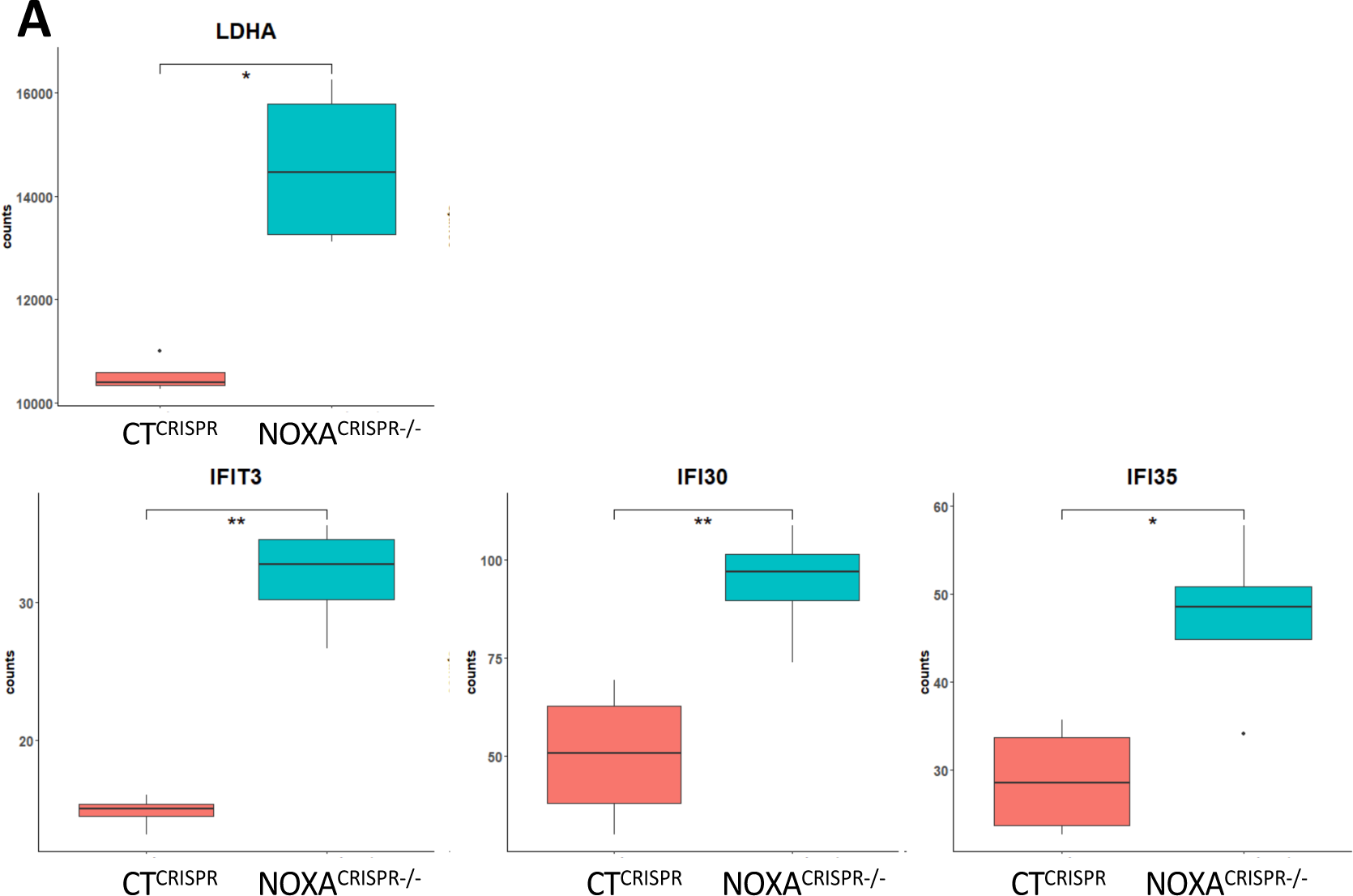
Targeted comparative proteomic analysis of secretomes. **A**: LDHA and **B**: IFIT3, IFI30 and IFI35 comparative release between taxol-treated CT^CRISPR^ and taxol-treated NOXA^CRISPR-/-^ MDA-MB-468 cells.

**Supplementary figure 3:**
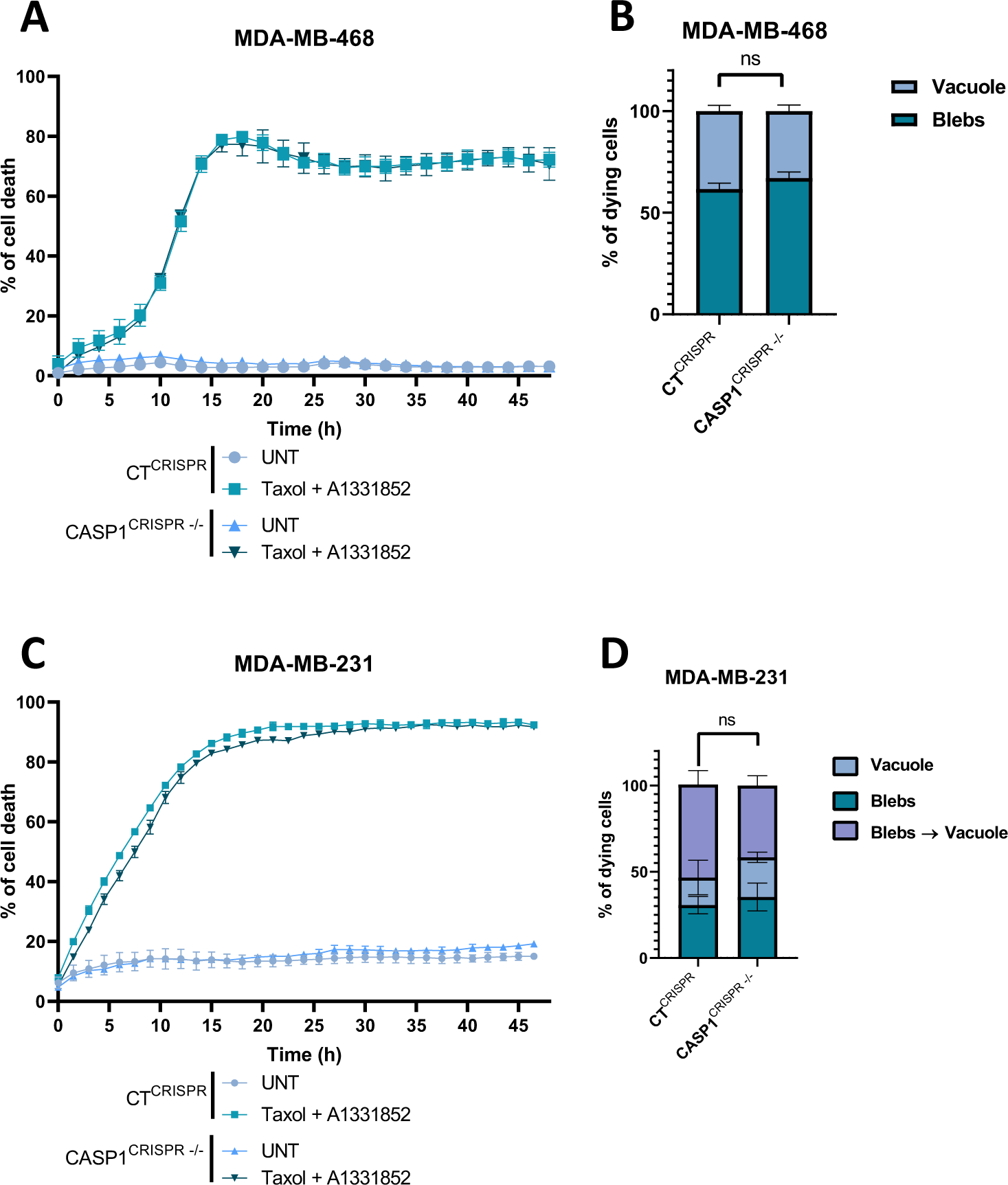
Cell death process was not modified in CASP1^CRISPR-/-^ compared to CT^CRISPR^ cells. **A, C**: Kinetics of single cell death analysis in not treated (UNT) or treated by taxol + A1331852 in combination during 48h, in CT^CRISPR^ and CASP1^CRISPR-/-^ MDA-MB-468 and MDA-MB-231 cells, respectively. **B,D:** % of cells dying either with blebs or with a unique vacuole after a 48h-exposure to combined treatment in CT^CRISPR^ and CASP1^CRISPR-/-^ MDA-MB-468 and MDA-MB-231 cells, respectively.

